# Analysis of Skin Cancers from Xeroderma Pigmentosum Patients Reveals Heterogeneous UV-Induced Mutational Profiles Shaped by DNA Repair

**DOI:** 10.1101/2022.10.14.512263

**Authors:** Andrey A. Yurchenko, Fatemeh Rajabi, Tirzah Braz-Petta, Hiva Fassihi, Alan Lehmann, Chikako Nishigori, Ismael Padioleau, Konstantin Gunbin, Leonardo Panunzi, Fanny Morice-Picard, Pierre Laplante, Caroline Robert, Patricia L. Kannouche, Carlos F. M. Menck, Alain Sarasin, Sergey I. Nikolaev

## Abstract

Xeroderma pigmentosum (XP) is a genetic disorder caused by mutations in genes of the Nucleotide Excision Repair (NER) pathway (groups A-G) or in Translesion Synthesis (TLS) DNA polymerase η (V). XP is associated with an increased skin cancer risk, reaching, for some groups, several thousand-fold compared to the general population. Here, we analyzed 38 skin cancer genomes from five XP groups. We found that the activity of NER determines heterogeneity of the mutation rates across skin cancer genomes and that transcription-coupled NER extends beyond the gene boundaries reducing the intergenic mutation rate. Mutational profile in XP-V tumors and experiments with *POLH*-KO cell line revealed the role of polymerase η in the error-free bypass of (i) rare TpG and TpA DNA lesions, (ii) 3’ nucleotides in pyrimidine dimers, and (iii) TpT photodimers. Our study unravels the genetic basis of skin cancer risk in XP and provides insights into the mechanisms reducing UV-induced mutagenesis in the general population.

## INTRODUCTION

Xeroderma Pigmentosum (XP) is a group of eight rare hereditary recessive disorders caused by mutations in seven nucleotide excision repair (NER) pathway genes (groups A-G) or in the *POLH* gene coding the translesion synthesis (TLS) DNA polymerase η (XP-V)^1^. XP is characterized by up to a 10000-fold increased risk of non-melanoma skin cancers and 2000-fold increased risk of melanoma^2^. Moreover, epidemiological studies revealed a 34-fold increased risk of internal tumors in XP patients which was associated with characteristic mutation signature and accelerated accumulation of mutations^3,4^.

Nucleotide excision repair (NER) is the main pathway that removes bulky DNA lesions in the genome in an error-free manner^5^. NER can be initiated by two sub-pathways: global genome repair (GG-NER) and transcription-coupled repair (TC-NER), while the downstream mechanism of lesion removal is shared between the two and involves recruitment of the TFIIH complex and XPA, which unwind the DNA helix at the lesion site, and XPG and XPF-ERCC1, which excise the fragment containing the damaged nucleotides^6^. GG-NER operates genome-wide; it recognizes UV-induced bulky lesions with the XPE/DBB2 protein or helix distortions caused by those lesions with XPC protein. TC-NER is initiated by lesion-stalled RNA polymerase II and operates mainly on the transcribed strand of active genes.

The photoproducts which have not been removed by NER may block the progression of replicative polymerases during DNA replication. The TLS polymerase η is a DNA polymerase that bypasses the UV-induced photoproducts, thus preventing replication fork stalling^7,8^.

Skin cancer predisposition among XP groups is highly heterogenous, and there is an inverse relationship between the level of sunburn sensitivity and skin cancer incidence between the groups^9^. The most skin cancer-prone groups are XP-C and XP-E, with impaired GG-NER, and XP-V - with the deficiency of polymerase η^9^. Other XP groups with the deficiency in both GG-NER and TC-NER are also associated with considerable skin cancer risk and demonstrate a high level of sunburn sensitivity and neurological symptoms^10^. A current model posits that UV exposure in the context of TC-NER deficiency may cause an impairment of transcription and result in decreased cell fitness, whereas defective GG-NER results in tumor-proneness^11^.

The process of UV-induced mutagenesis depends on several major factors, including DNA lesion generation, removal by NER, and bypass by TLS polymerases. Skin cancers from XP groups differ from each other and sporadic skin cancers by the ability to repair or bypass DNA lesions, but not by the sources of DNA damage. Thus, analysis of XP tumors with defects in GG-NER alone, both TC- and GG-NER, or TLS, enables the disentanglement of the contribution of those components to mutagenesis in a natural physiological system. Moreover, an extreme skin cancer susceptibility in XP patients points to vulnerabilities in the mechanisms of protection from excessive mutation accumulation in normal skin cells.

In this study, we assembled a unique collection of 38 skin cancers from 5 xeroderma pigmentosum groups (XP-A, C, D, E, V). We used Whole Genome Sequencing (WGS) to study the role of defects in the major components of NER and translesion synthesis on tumor mutation burden, mutation profiles, genomic landscape, and protein-damaging effects of mutagenesis in human skin cancers.

## RESULTS

### Samples and clinical characteristics

We collected and sequenced genomes of 33 skin cancers from 21 patients representing 5 out of 8 xeroderma pigmentosum (XP) groups (3 XP-A, 4 XP-C, 2 XP-D, 10 XP-E, and 14 XP-V tumors; **Supplementary Table 1**). Causative homozygous (n=12) or compound heterozygous (n=8) germline variants were identified in 20 patients, 13 of which had known causative germline mutations (**Supplementary Table 1**), while the 7 others – had novel germline mutations compatible with the diagnosis. The mean tumor purity and sequence coverage were 41% and 40X (30X for normal tissue), respectively. Additionally, we sequenced genomes of 6 sporadic cutaneous squamous cell carcinoma samples (SCC). This newly generated data was combined with WGS data from four previously published XP-C tumors^12,13^, one XP-D^14^, as well as 25 sporadic cutaneous Squamous Cell Carcinomas (SCC)^12,15,16^, 8 Basal Cell Carcinomas^15^ (BCC) and 113 Melanomas^17^ (MEL) from individuals not affected with XP. The resulting cohort of XP tumors included 17 BCCs, 15 SCCs, five melanomas, and one angiosarcoma. The mean age at biopsy in XP-cohort was 33 years old (ranging from 25 years old in the XP-C group to 48 years old in the XP-V group) while in sporadic skin cancer group it was 65 years old (**Table 1, Supplementary Table 1**).

**Table 1.** The studied dataset of XP and sporadic skin cancers with WGS data.

| Group | Tumors (n) | Patients (n) | Mean age at biopsy (years) | cSCC (n) | BCC (n) | Melanoma (n) |
| --- | --- | --- | --- | --- | --- | --- |
| XP-E | 10 | 3 | 28 | 7 | 2 | 1 |
| XP-C | 8* | 8 | 25 | 5 | 1 | 1 |
| XP-A | 3 | 1 | 32 | 1 | 2 | - |
| XP-D | 3 | 3 | 30 | 2 | 1 | - |
| XP-V | 14 | 9 | 48 | - | 11 | 3 |
| Sporadic SCC | 31 | 31 | 73 | 31 | - | - |
| Sporadic BCC | 8 | 8 | 66 | - | 8 | - |
| Sporadic MEL | 113 | 113 | 57 | - | - | 113 |

\*- one XP-C patient with angiosarcoma

### XP groups demonstrate different mutation burden and mutation profiles

We assessed the Tumor Mutation Burden (TMB) and mutation profiles of skin cancer genomes from 5 sequenced XP groups and compared them with the three types of sporadic skin cancer including BCC, SCC and MEL (**Fig. 1a**). The mean TMB of single base substitutions (SBS) was significantly higher in 3 XP groups: XP-E (350 mut/Mb, *P* = 0.0241), XP-V (248 mut/Mb, *P*=0.0014) and XP-C skin cancers (162 mut/Mb, *P*=0.0220), than the dataset weighted average (130 mut/Mb, global *P* < 2.2e-16; Kruskal–Wallis H test; **Fig. 1a**). We also observed a striking difference in the TMB and the proportion of CC>TT tandem base substitutions (DBS) characteristic of UV-induced mutagenesis between the different XP groups and sporadic cancers (**Fig. 1a**). The highest proportion of CC>TT DBS from UV-induced SBS in pyrimidine dimers (C>T in Yp<u>C</u> or <u>C</u>pY contexts; Y denotes a pyrimidine) was observed in XP-C and XP-D tumors (0.2 and 0.17, respectively), which was 6 times higher than in sporadic skin cancers (0.03, *P* = 4.7e-08, Mann–Whitney U test, two-sided).

**Figure 1.**
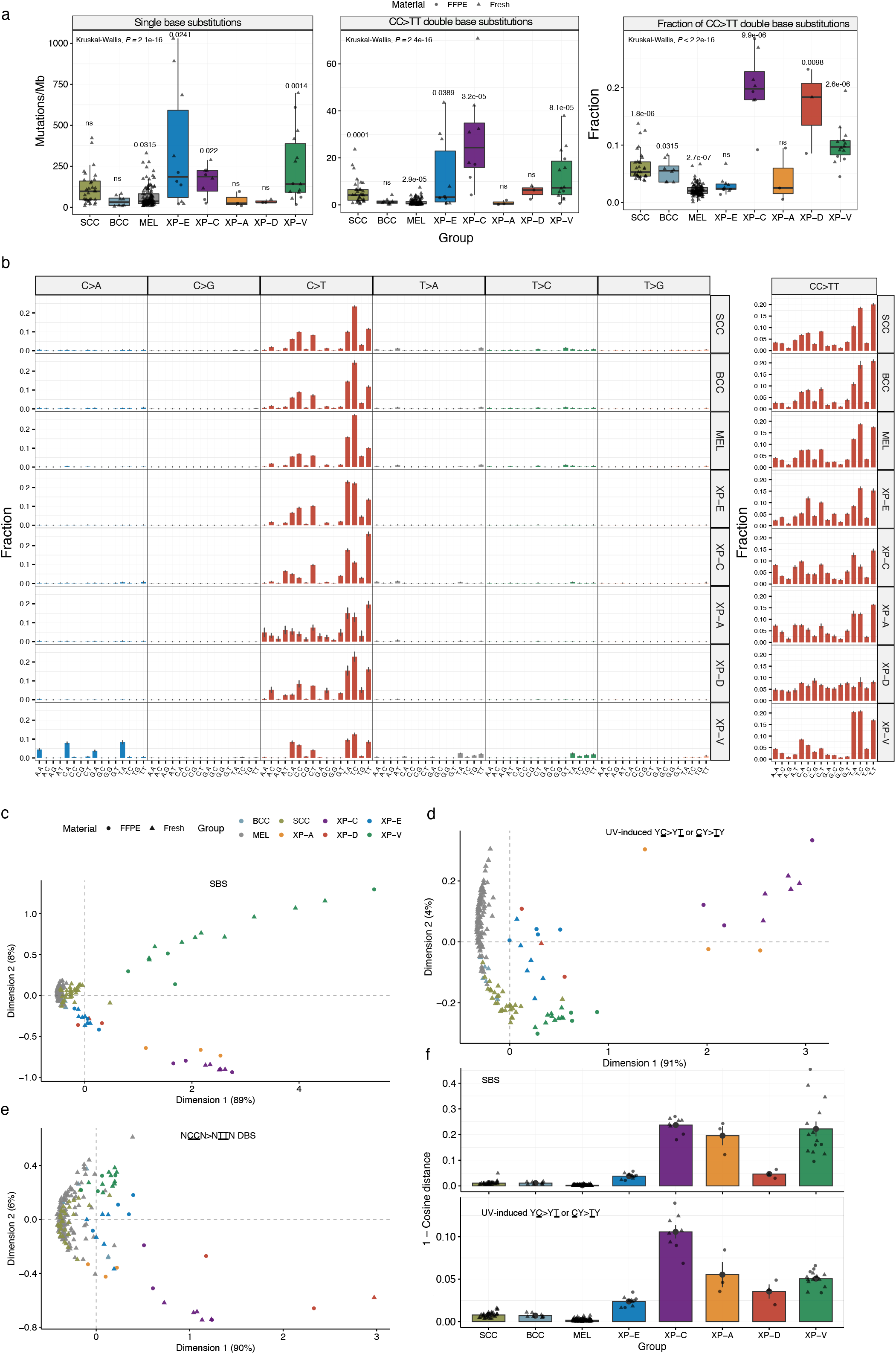
Mutation landscape of the studied cancers. **a** Tumor mutation burden of single base substitutions (SBS; left panel), double base substitutions (CC>TT; middle panel) per group and a proportion of CC>TT DBS relative to C>T SBS in pyrimidine context (right panel). *P*-values from nonparametric ANOVA are indicated. **b** Trinucleotide-context mutation profiles of SBS (left panel) and tetranucleotide-context mutation profiles of CC>TT DBS (right panel) per group. **c** Multidimensional scaling (MDS) plot based on the Cosine similarity distance between the SBS trinucleotide-context mutation profiles of the samples. **d** MDS plot based on the Cosine similarity distance between the trinucleotide-context mutation profiles of the samples using only C>T mutations with an adjacent pyrimidine (Y<u>C</u>>Y<u>T</u> or <u>C</u>Y><u>T</u>Y), the typical UV mutation context. **e** MDS plot based on the Cosine similarity distance between the tetranucleotide-context mutation profiles of the samples using only CC>TT double base substitutions. **f** Mean Cosine dissimilarity (1-Cosine distance) between original and reconstructed trinucleotide-context mutation profiles using only SBS7a/b/c/d COSMIC mutation signatures for all SBS (upper panel) and C>T mutations with adjacent pyrimidine only (lower panel).

The mutation profiles of skin cancers in all XP groups were dominated by C>T substitutions at pyrimidine dimers, as also found in sporadic skin cancers. However, some XP groups demonstrated marked differences for C>T mutations in specific contexts, such as enrichment at T<u>C</u>A in XP-E, T<u>C</u>W in XP-C, or N<u>C</u>Y in XP-D (where W denotes A or T; N: A, C, G, or T; Y: C or T). Moreover, in XP-V skin cancers, we report abundant mutations, namely C:G>A:T, T:A>A:T, and T:A>C:G, which were not previously seen to a significant degree in skin cancer (**Fig. 1b, Supplementary Fig. 1**). XP tumors formed clusters by XP group, which were non-overlapping with the cluster of sporadic skin cancers based on SBS mutation profiles and multidimensional scaling analysis (MDS; **Fig. 1c**,**d**,**e**). XP-V, XP-C, and XP-A clusters were located distantly, while the XP-E / XP-D cluster was closer to the cluster of sporadic skin cancers.

Among 78 COSMIC mutation signatures^18^ (v3.2) extracted from the pan-cancer dataset, four mutation signatures (SBS7a/b/c/d) are associated with UV irradiation. We investigated whether these signatures could explain the observed mutation profiles in XP skin cancer with an accuracy comparable to sporadic skin cancers. For that, we compared observed and reconstructed mutation profiles for each sample in our cohort. The mean Cosine dissimilarity distance was small for sporadic skin cancers (0.004) but increased drastically for all the XP groups (0.16) and particularly for XP-C (0.237), XP-A (0.1957), and XP-V (0.222, **Fig. 1f**).

### Nucleotide excision repair efficiency determines mutation load distribution along the genome

Strong heterogeneity in the mutation rate across the genome is an important fundamental feature of mutagenesis, which has several clinical implications, for example, the discovery of cancer driver genes. We investigated the distribution of typical UV mutations (Y<u>C</u>>Y<u>T</u> or <u>C</u>Y><u>T</u>Y) in XP and sporadic skin cancers in relation to replication timing (RT), active and inactive topologically-associated domains (TAD), and markers of chromatin states. These analyses revealed a major role for NER in shaping the heterogeneity of local rates of UV-induced mutations across the genome. A maximal 5.2-fold difference was observed between the earliest and the latest replicating bins in sporadic skin cancers (average for BCC, cSCC, and MEL) with a monotonal decrease of mutation load from late to early replicating genomic regions (**Fig. 2a**). This effect was much weaker in GG-NER deficient XP-C genomes (2.4-fold) and almost disappeared in GG-NER and TC-NER deficient XP-A (1.5-fold) and XP-D (0.99-fold) genomes (**Fig. 2a**). Interestingly, the distribution of UV-induced SBS by RT in XP-E and XP-V genomes was not very different from sporadic skin cancer genomes, 4.6-fold and 5.4-fold, respectively.

**Figure 2.**
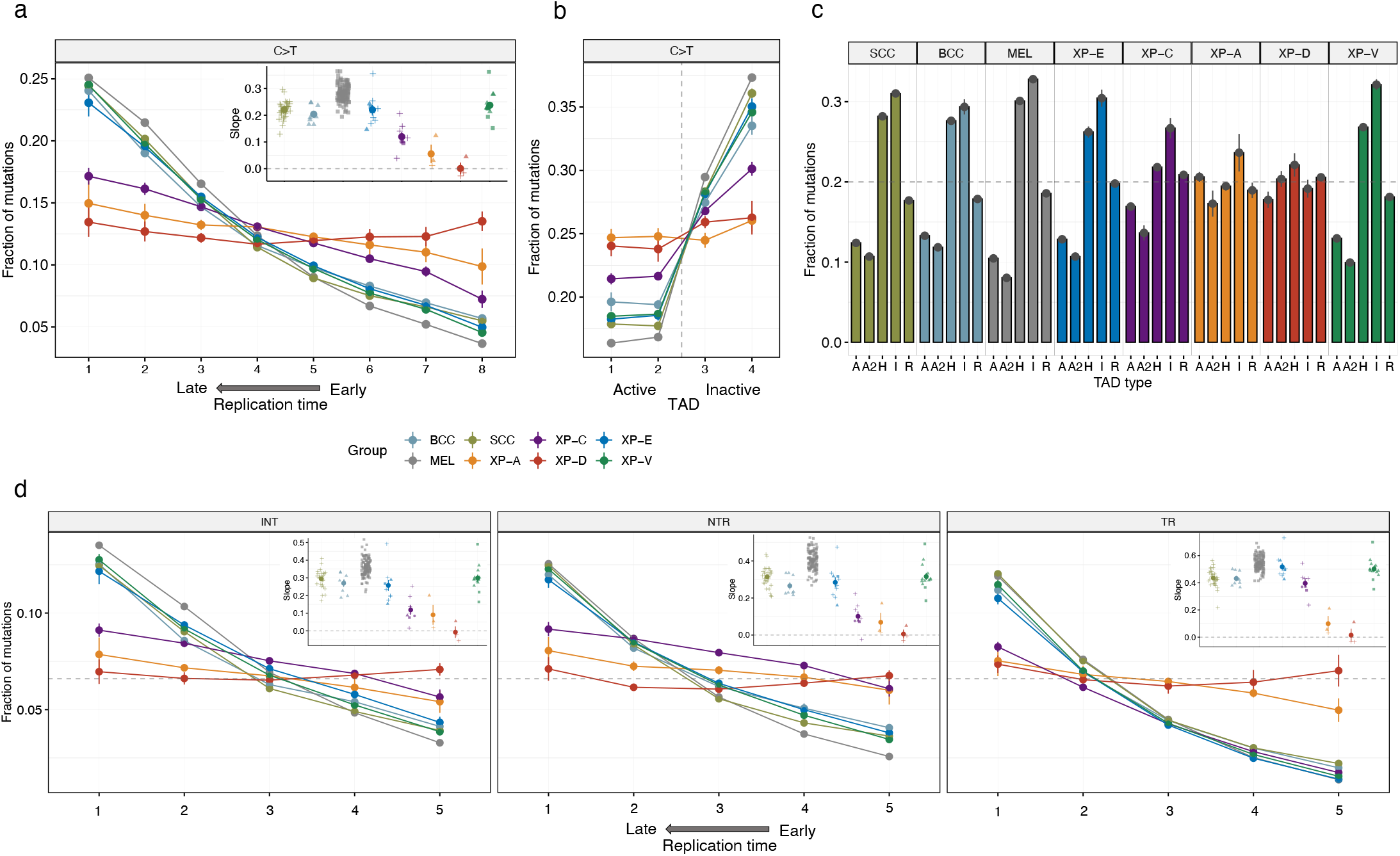
Genomic topography of mutagenesis in the skin cancers. **a** Fraction of C>T mutations from pyrimidine dimers in genomic regions grouped in 8 equal size bins by replication timing (RT) for XP groups and sporadic skin cancers. The box contains the slope values from linear regressions across 8 RT bins. **b** Fraction of C>T mutations from pyrimidine dimers per group in 1Mb regions centered at the boundary between active and inactive topologically-associated domains (split into two bins each). **c** Fraction of C>T mutations from pyrimidine dimers per group across different chromatin states (R - repressed, A and A2 - active, H - heterochromatin, I - inactive). **d** Fractions of C>T mutations from pyrimidine dimers in intergenic regions (left panel), on the untranscribed (middle panel) and transcribed (right panel) DNA strands of gene regions grouped in 5 equal size bins by replication timing (RT) for XP groups and sporadic skin cancers. The boxes contain the slope values from linear regressions across 5 RT bins. I: intergenic regions; NTR: untranscribed strand of genes; TR: transcribed strand of genes.

It has been recently shown that TAD boundaries between active and inactive chromatin domains strongly delineate the transition between regions with low and high mutation load in different human cancers^19^. Indeed, in our cohort, we found a 2.2-fold difference in mutation load between active and inactive TADs in sporadic cancers, but it was noticeably decreased in XP-C (1.4-fold) cancers and was virtually absent in XP-A (1.05-fold) and XP-D (1.09-fold; **Fig. 2b**). Similarly, the mutation load in XP-A and XP-D tumors was independent of chromatin states, the XP-C group demonstrated a mild dependence, while the XP-E and the XP-V groups were not different from sporadic cancers (**Fig. 2c**).

CPD and 6-4PP DNA lesions occur on pyrimidine bases, which enabled us to identify the strand on which the lesion underlying a UV-induced mutation occurred. In order to separately investigate the genomic targets of GG-NER and TC-NER, we split the genome into intergenic, transcribed, and untranscribed strands of genic regions. A strong decrease mutation rate in the early RT regions in groups proficient in GG-NER (sporadic cancers and XP-V), and surprisingly in GG-NER deficient XP-E, was observed in intergenic regions and untranscribed strands of genes. Whereas XP-A, XP-D, and XP-C groups, which lack GG-NER, had flat slopes compatible with the lack of repair in the open chromatin of early RT regions (**Fig. 2d, Supplementary Fig. 2)**.

### Transcriptional bias is different between the XP groups

TC-NER removes UV-induced bulky DNA lesions on the transcribed strand of expressed genes more efficiently than GG-NER on the untranscribed strand resulting in a decrease of mutations on the transcribed versus untranscribed strand, a phenomenon called transcriptional bias (TRB)^20^. In skin tumors with proficient NER, the TRB ranged between 1.3 and 1.6-fold for sporadic cancers and was 1.7 in XP-V. In the GG-NER-deficient TC-NER proficient groups, TRB was particularly high, ranging between 1.77-fold (XP-E) and 2.42-fold (XP-C), which is compatible with defects in the repair of the untranscribed strand. In contrast, in XP-A and XP-D groups with defects of both TC-NER and GG-NER TRB was minimal or absent: 1.17-fold and 0.97-fold, respectively (**Fig. 3a**).

**Figure 3.**
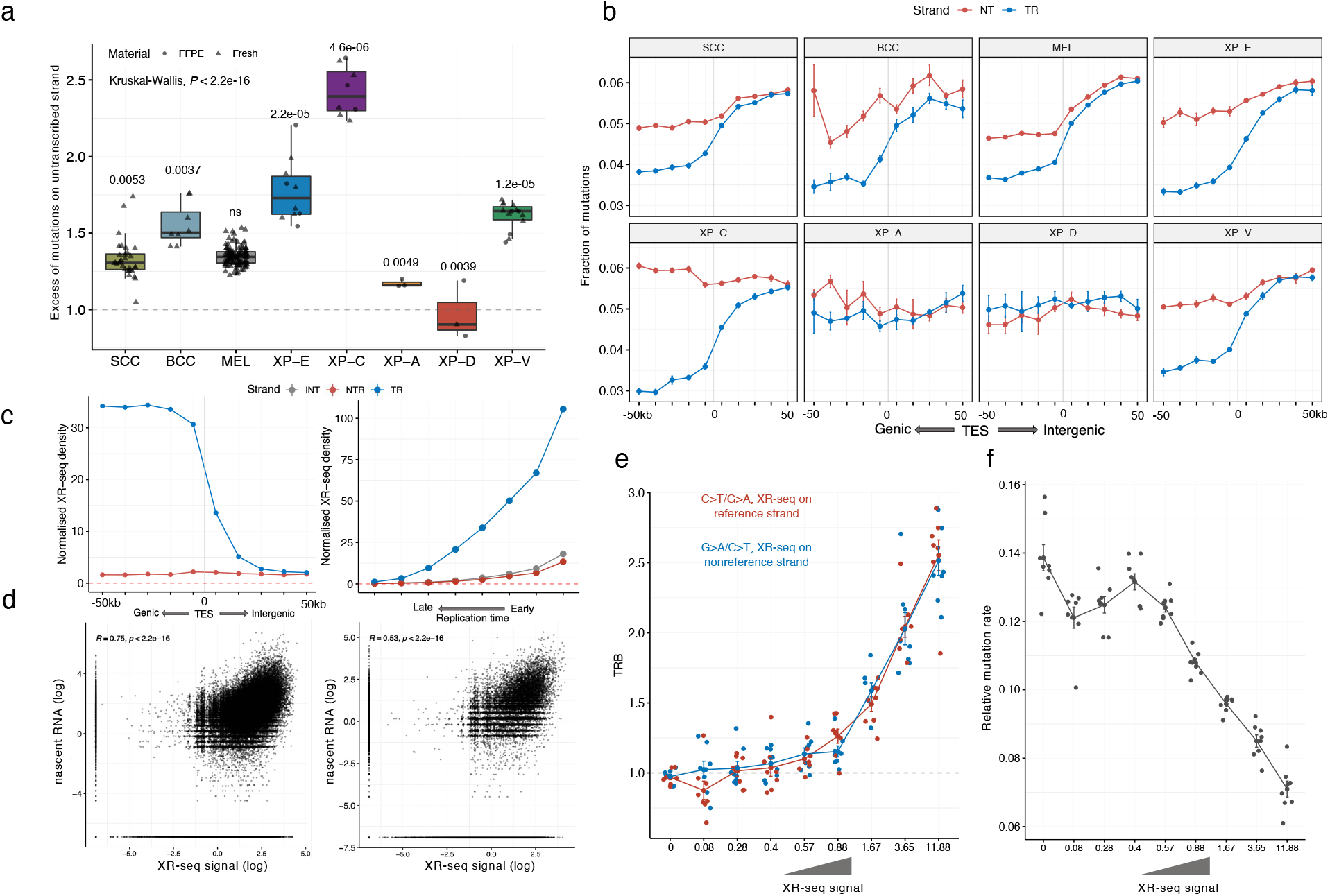
TC-NER activity behind transcription end sites (TES) of genes. **a** The transcriptional bias (TRB) per group (ratio between untranscribed and transcribed strand) for C>T mutations with adjacent pyrimidines for XP groups and sporadic skin cancers. *P*-values from nonparametric ANOVA are indicated **b** Fractions of C>T mutations with adjacent pyrimidines separated by strands in the TES-centered 100kb region (binned by 10kb intervals). **c** DNA context-normalized XR-seq density from XP-C cell line on untranscribed and transcribed gene strands in the TES-centered 100kb region (binned by 10kb intervals; left panel). DNA context-normalized XR-seq density from XP-C cell line by replication timing for the transcribed and untranscribed DNA strands of genes and intergenic regions. I: intergenic regions; NTR: untranscribed strand of genes; TR: transcribed strand of genes (right panel). **d** Correlation between XR-seq intensity from XP-C cell line and nascent RNA-seq for genic regions (left panel) and intergenic regions 50kb downstream of TES (right panel). **e** Transcriptional bias of C:G>T:A mutations on intergenic regions of XP-C tumors depending on the XR-seq intensity of XP-C cell line. **f** Relative mutation rate of C:G>T:A mutations in intergenic regions of XP-C tumors depending on the XR-seq intensity in XP-C cell line.

### TC-NER removes DNA lesions downstream of genes and influences intergenic mutation load

Since early RT regions are particularly gene-rich^21^, we hypothesized that in GG-NER deficient XP groups, decreased mutation load in early RT intergenic regions might be associated with the TC-NER activity beyond gene boundaries. Indeed, in GG-NER deficient XP-C tumors, we revealed a significant TRB up to 50kb downstream of the furthest annotated transcriptional end sites (TES) of genes with decreased mutation frequency on the transcribed strand of nearby genes (**Fig. 3b, Supplementary Fig. 3**). The same effect was observed in XP-E and even in NER proficient skin cancers although with a lower magnitude. As expected, we did not observe TRB downstream of genes in XP-A and XP-D samples being deficient for both TC-NER and GG-NER (**Fig. 3b, Supplementary Fig. 3**). To validate TC-NER activity downstream of gene TES, we used previously published XR-seq data from XPC-deficient cell lines. It is expected that in XPC-deficient cells, XR-seq data, representing the sequencing of lesion-containing DNA fragments excised by NER^22^, is produced exclusively by TC-NER. An XR-seq signal was observed up to 50kb downstream of TES on a transcribed strand of a nearby gene, mirroring mutation asymmetry in the same regions in XP-C tumors (**Fig. 3c**) and was well correlated with the transcriptional intensity of nascent RNA, which was retrieved from an independent study ^23^ (**Fig. 3d**). This suggests that, in some cases, the RNA polymerase might continue transcription after TES and recruit TC-NER at lesion sites. We identified XR-seq signal in 21% of the cumulative length of intergenic regions and 14% - of untranscribed strands of genes in XPC-deficient cell line^22^, suggesting ubiquitous extended TC-NER activity. Analysis of transcriptional bias and relative mutation rate in intergenic regions of XP-C tumors (**Fig. 3e, f**) revealed strong dependence on the intensity of XR-seq outside the annotated genic regions. This extended TC-NER activity outside of the transcribed strand of genes is especially strong in early replicating regions with a high density of active genes (**Fig. 3c**). It may explain the decrease of the mutation density in intergenic regions and on the untranscribed strands of genes in early replicating genomic regions of GG-NER deficient XP-C samples (**Fig. 2d, Supplementary Fig. 2**).

### XP-E demonstrates reduced GG-NER activity

The sensors of UV-induced DNA lesions in GG-NER, XPC, and DDB2 (XPE) are thought to work in tandem when DDB2 binds directly to a lesion and facilitates recruitment of XPC, which in turn initializes the repair process with the TFIIH complex^24^. We next decided to compare the features of UV-induced mutagenesis in XP-E resulting from the loss of DDB2 with XP-C and sporadic tumors.

MDS plot based on SBS revealed three different well delineated clusters of XP-C, XP-E, and sporadic tumors (**Fig. 4a**). At the same time, the proportion of CC>TT DBS was much increased in XP-C (0.21) versus sporadic cSCC (0.064), but significantly decreased in XP-E cSCC (0.034, *P* = 0.0003; Mann–Whitney U test, two-sided), confirming qualitative differences of mutagenesis in XP-E. Unlike XP-C, the distribution of the mutational load in intergenic and untranscribed strand gene regions by RT in XP-E was very close to that of sporadic cSCC, suggesting that repair in early RT regions was functional in XP-E (**Fig. 2d, Supplementary Fig. 2**). Similarly, the MDS based on the local mutation load in 2684 1Mb-long intervals along the genome, revealed no difference between XP-E and sporadic samples, while XP-C formed a separate cluster irrespective the tumor type (**Fig. 4b**).

**Figure 4.**
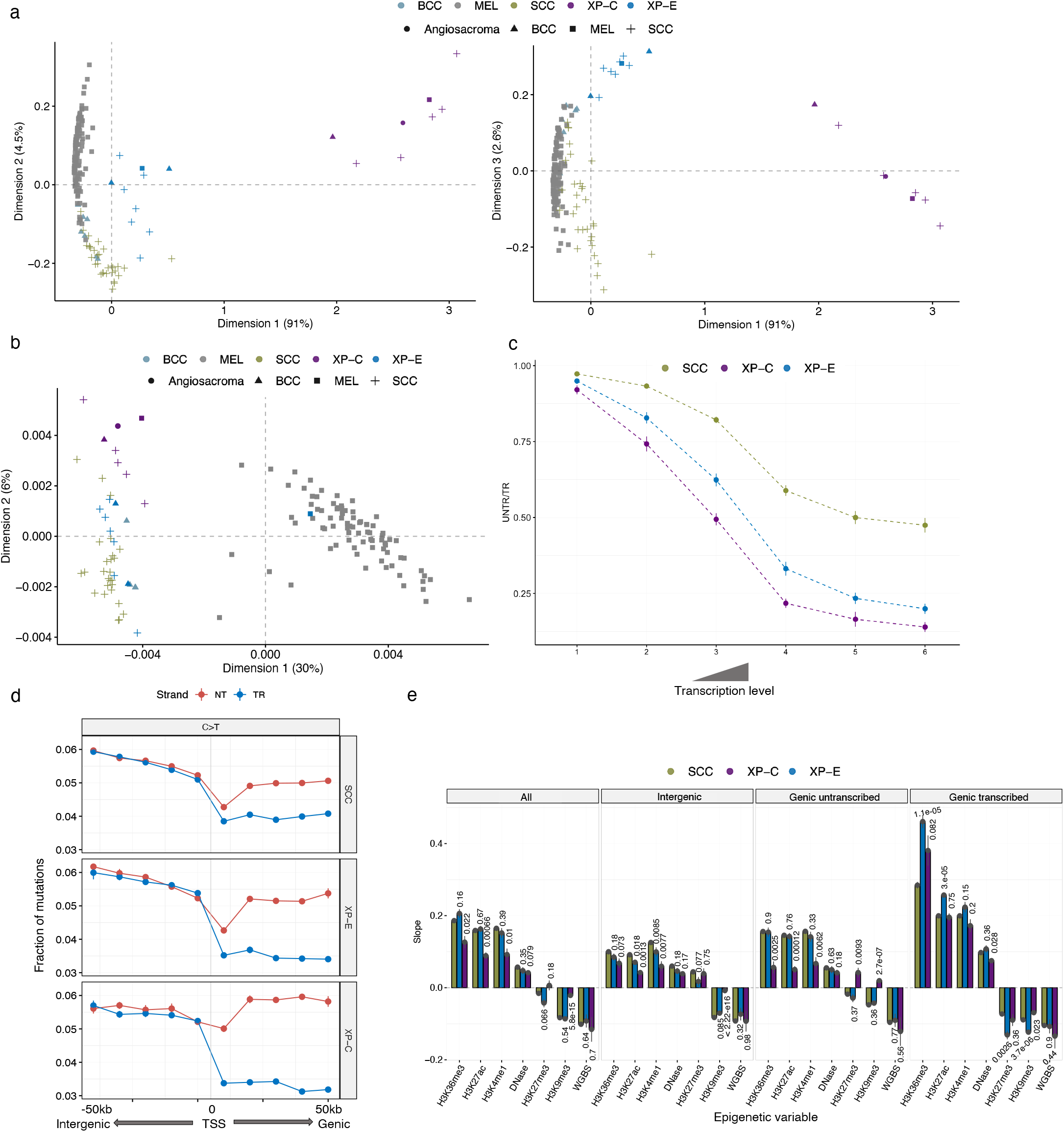
Comparison of genomic mutagenesis between sporadic cancers, XP-E and XP-C groups. **a** Multidimensional scaling (MDS) plot based on the Cosine similarity distance between the SBS trinucleotide-context mutation profiles of the samples (Dimensions 1 and 2 - left panel, Dimensions 1 and 3 – right panel). **b** PCA plot based on the density of mutations in 2684 1Mb-long windows along the genome (only for samples with more than 50k mutations belong to sporadic, XP-C and XP-E groups). **c** The transcriptional bias (TRB; ratio between untranscribed and transcribed strand mutation number) for C>T mutations from pyrimidine dimers in genes grouped in 6 bins by gene expression level. Only cSCC tumors were used for XP-C and XP-E groups **d** Fractions of C>T mutations from pyrimidine dimers separated by strands in the TSS-centered 100kb region (binned by 10kb intervals). **e** The slope values from linear regressions across C>T mutations from pyrimidine dimers over binned epigenetic features for the whole genome (left panel), intergenic regions (left middle panel), untranscribed (right middle panel), and transcribed (right panel) strands of genes separately (only cSCC from sporadic, XP-E and XP-C groups were used in the analysis). *P*-values based on the Student’s t-test pairwise comparisons between sporadic cSCC and cSCC from XP-C or XP-E groups are indicated.

The XP-E group demonstrated a strong TRB (1.77-fold), which was intermediate between sporadic cSCC (1.33) and the XP-C group (2.47) (**Fig. 3a, Fig. 4c**,**d**). Given that TC-NER is functional in XP-E, XP-C, and sporadic samples, and assuming that GG-NER is fully abrogated in XP-C, we can estimate the relative efficiency of GG-NER in XP-E tumors. Providing all else is equal, GG-NER is 64% less efficient in XP-E than in sporadic cancers.

To provide a more detailed view of the mutation difference between XP-E, XP-C, and sporadic tumors, we compared the association of mutation load in each group with the core epigenetic marks from primary keratinocyte cell line^25^ using only cSCC samples (**Fig. 4e**). Unlike XP-C, XP-E tumors did not show strong and significant differences from sporadic cSCC in the dependence of mutagenesis on the majority of epigenetic covariates except for the histone modification marks H3K36me3, H3K27ac and H3K9me3 on the transcribed strand of gene regions (**Fig. 4e**).

Taking these observations together, we can speculate that in XP-E tumors, there is a residual activity of GG-NER associated with the ability of XPC to find a fraction of DNA lesions and initiate NER. This correlates with the clinical observation that XP-E patients develop less and later skin tumors than XP-C patients.

### Polymerase η deficiency causes a unique mutation profile in skin cancers

The analysis of XP-V skin cancers revealed that an average of 27% (15-42%) of SBS were represented by C:G>A:T mutations with a highly specific 3-nt context (N<u>C</u>A) and a strong and homogeneous TRB (**Fig. 5a, Fig. 1b, Supplementary Fig. 1**). Similar mutation contexts and a TRB was observed for a part of T:A>A:T mutations, which represented 8.7% of SBS. In sporadic skin cancers, C:G>A:T and T:A>A:T mutations represented only 2.5% and 4.6%, respectively, and had different broad 3-nt contexts without a strong TRB (**Fig. 1b, Supplementary Fig. 1**). Enrichment of these types of mutations in XP-V suggests that they might originate from lesions that are bypassed by polymerase η in an error-free manner in sporadic skin cancer, but XP-V cells have to use an alternative polymerase(s) to bypass these lesions.

**Figure 5.**
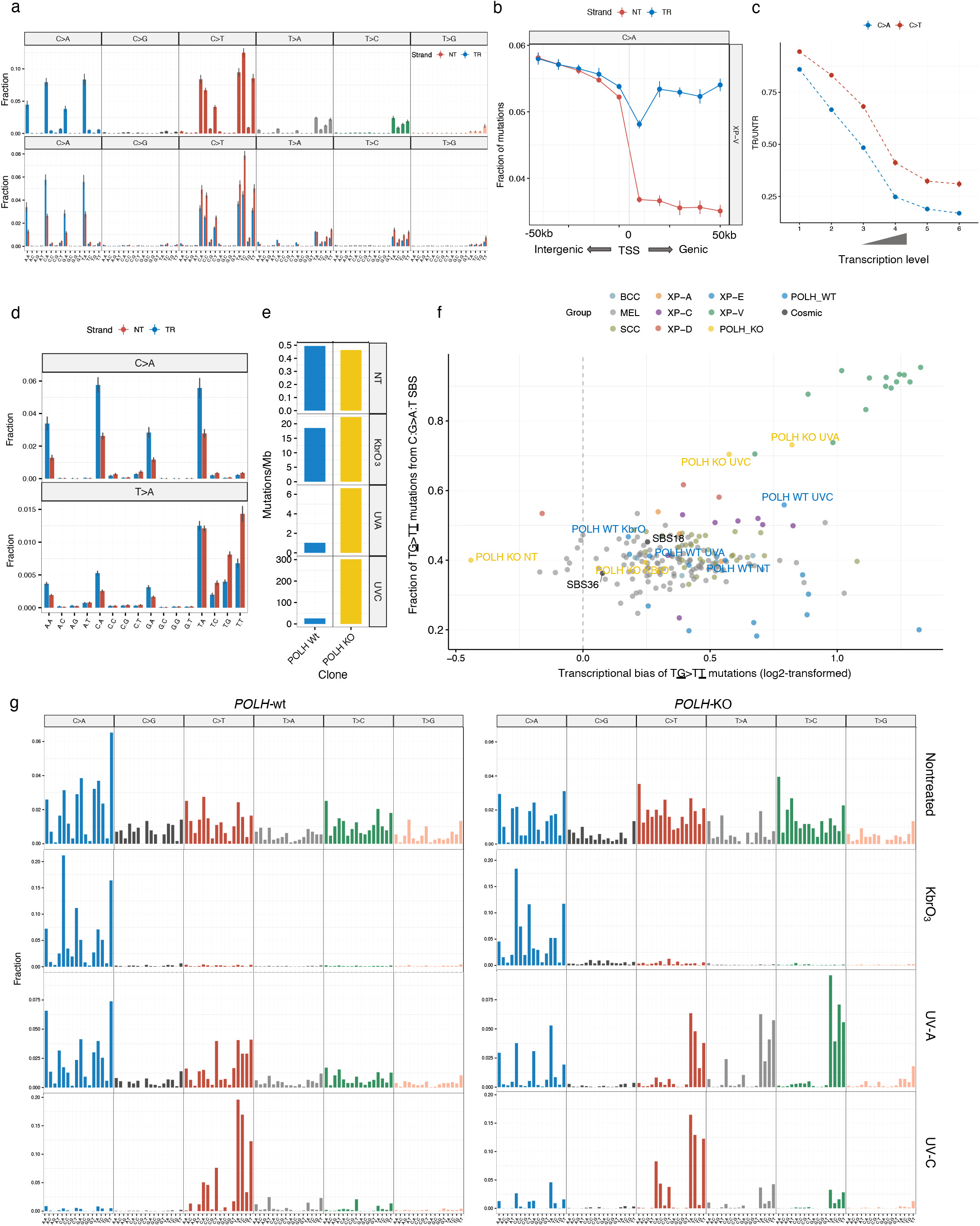
Mutation profiles of XP-V skin cancers and *POLH*-KO clones. **a** Trinucleotide-context mutation profile of genomic SBS (upper panel) and genic SBS (lower panel) separated by transcribed (TR) and untranscribed (NT) strands in XP-V tumors. **b** Fractions of C>A mutations separated by gene strands in the TSS-centered 100kb region of XP-V tumors (binned by 10kb intervals). Blue – transcribed strand for mutations from purines and untranscribed strand for mutations from pyrimidines; red - untranscribed strand for mutations from purines and transcribed strand for mutations from pyrimidines **c** The transcriptional bias (ratio between transcribed and untranscribed strand) for C>A and C>T mutations per bin of gene expression level (only XP-V samples represented by SCC and BCC). **d** Trinucleotide-context mutation profiles of SBS separated by strands in XP-V tumors for C>A and T>A mutations. **e** Mutations per megabase in the *POLH* wt and *POLH*-KO clones in nontreated cells (NT), treated with KbrO_3_, UV-A and UV-C. **f** Mutational specificity of the T<u>G</u>>T<u>T</u> mutations in XP-V tumors and *POLH*-KO UV-A- and UV-C-treated cell lines. X-axis: log2-transformed transcriptional bias of the T<u>G</u>>T<u>T</u> mutations per genome. Y-axis: Fraction of the mutations in the T<u>G</u>>T<u>T</u> context from the total number of C:G>A:T substitutions per genome. *POLH*-KO and *POLH*-wt clones are specifically indicated with their corresponding treatment (KbrO_3_, UV-A and UV-C) as well as COSMIC SBS18 and SBS36 mutational signatures associated with oxidative DNA damage (black dots). **g** Mutation profiles of the *POLH*-wt and *POLH*-KO clones for nontreated cells (NT), treated with KbrO_3_, UV-A and UV-C.

The direction of TRB for these types of mutations indicates a decrease in mutations from lesions involving purines on the transcribed strand (**Fig. 5a**). Furthermore, comparison of C:G>A:T mutation frequencies on the transcribed and untranscribed strands with the proximal 5’ intergenic regions confirmed that TRB is indeed associated with a decrease of C:G>A:T mutations on the transcribed strand (**Fig. 5b**). This suggests that mutations occur due to lesions involving purines, which are NER substrates and are effectively repaired by TC-NER on the transcribed strand (**Fig. 5b**). Interestingly, C:G>A:T mutations had stronger TRB than Y<u>C</u>>Y<u>T</u> or <u>C</u>Y><u>T</u>Y UV-induced mutations in all bins of genes grouped by the expression level (**Fig. 5c**). This observation might indicate that those lesions produce a smaller helix distortion and are less visible to GG-NER than UV-induced pyrimidine lesions.

C:G>A:T and T:A>A:T mutations occurred in a very specific dinucleotide context, where a purine is always preceded by a thymine base (T<u>A</u>/<u>G</u> > T<u>T</u>), suggesting that causative DNA lesions might be thymine-purine dimers (**Fig. 5d**). The number of mutations in a T<u>G</u> context was strongly correlated with the number of mutations in a T<u>A</u> context (R=0.98; **Supplementary Fig. 4**) in our XP-V skin cancer cohort suggesting coordinated mutation processes.

To assess the possibility that these lesions were generated directly or indirectly due to UV-irradiation, we measured a Pearson correlation of T<u>G</u> > T<u>T</u> or T<u>A</u> > T<u>T</u> mutations with typical UV-induced (Y<u>C</u>>Y<u>T</u> or <u>C</u>Y><u>T</u>Y) mutations and observed strong correlations in both cases, *R*=0.78 (*P* = 0.001) and R=0.99 (*P* = 1e-10), respectively (**Supplementary Fig. 4**).

To further understand the nature of T<u>G</u> > T<u>T</u> and T<u>A</u> > T<u>T</u> mutations we established a *POLH* knock out of the RPE-1 *TP53*-KO cell line and sequenced whole genomes of the *POLH* wt and *POLH*-KO clones both without treatment and with treatment with KbrO3 (to induce reactive oxygen species), UV-A and UV-C (**Supplementary Fig. 5)**. There were no major differences in the number of mutations and mutational profiles between *POLH*-wt and *POLH*-KO for untreated cells and KbrO_3_-treated (**Fig. 5e**,**f**,**g**). UV-A and UV-C exposures greatly increased number of SBS in the *POLH*-KO cells (6.4 and 11.7-folds respectively) and dramatically changed the mutational profiles in comparison with *POLH*-wt clones (**Fig. 5e**,**f**,**g**). UV-A-treated *POLH*-KO clone had 15% of T<u>G</u> > TT mutations and 12% of the T<u>A</u> > T<u>T</u> mutations with specific XP-V context and strong transcriptional bias while in the UV-C-treated clone these percentages were 10% and 4% respectively (**Fig. 5f**,**g**). UV-treated *POLH*-KO cells demonstrated a distinct pattern of TG>VT DBS substitutions (V – A, C or G). Interestingly, a similar DBS pattern was also visible in XP-V tumors (**Supplementary Fig. 6**).

Another feature of the XP-V skin cancer profile was the presence of 15% (range 11% - 23%) of mutations originating from TT pyrimidine dimers. Such mutations are very rare in sporadic cancer (4.8%) because TT pyrimidine dimers are bypassed by polymerase η in a relatively error-free manner. Two predominant types of mutations at TT were T<u>T</u>>T<u>A</u> and T<u>T</u>>T<u>C,</u> and they, as expected for mutations from pyrimidine lesions, demonstrated strong TRB and were correlated with the typical UV-induced Y<u>C</u>>Y<u>T</u> or <u>C</u>Y><u>T</u>Y mutations (**Fig. 5a, Supplementary Fig. 4)**. Interestingly, UV-A treated *POLH*-KO cells harbored 52% of T>A and T>C mutations, while in case of treatment with UV-C, it was only 22% (**Fig. 5g**).

### In the absence of polymerase η, error-prone bypass of 3’ nucleotides in pyrimidine dimers shapes the mutation profile of XP-V tumors

The 3-nt context of C>T substitutions in XP-V skin cancers differed from sporadic skin cancers and other XP groups (**Fig. 1b**,**d**). Previously it was shown that in the absence of polymerase η, the bypass of CPD photoproducts can be performed in two steps by two TLS polymerases, one of which inserts a first nucleotide opposite to a 3’ nucleotide of the lesion (“inserter”), and then is replaced by another TLS polymerase, which performs the extension opposite to the 5’ nucleotide of the lesion (“extender”)^26^. We hypothesized that loss of polymerase η in skin cancer might change the probabilities of mutations at 3’ versus 5’ nucleotides in pyrimidine dimers and thereafter contribute to the observed differences of the mutation profiles for C>T SBS in XP-V versus sporadic skin cancer.

To test this hypothesis, we first estimated the relative number of mutations arising at 3’ and 5’ cytosines in the tetranucleotide ACCA, where we could unambiguously allocate a pyrimidine dimer (**Fig. 6a**). In sporadic skin cancers, the probabilities of mutations at 3’ and 5’ cytosines were similar, with only a slight increase of mutagenesis from the 3’C (55%), while in XP-V skin cancers 97% of the mutations were from the 3’C (**Fig. 6b**). This bias towards 3’ pyrimidine mutations was also much stronger in XP-V versus other groups of skin cancer for the CT, TC, and TT pyrimidine dimers. For example, AT<u>C</u>A > AT<u>T</u>A mutations were 9.17-fold more frequent than A<u>C</u>TA > A<u>T</u>TA mutations in XP-V than in the other groups (normalized to the corresponding 4-nt frequencies in the human genome). A similar effect was observed for T>A and T>C mutations in ATTA context (**Fig. 6c**).

**Figure 6.**
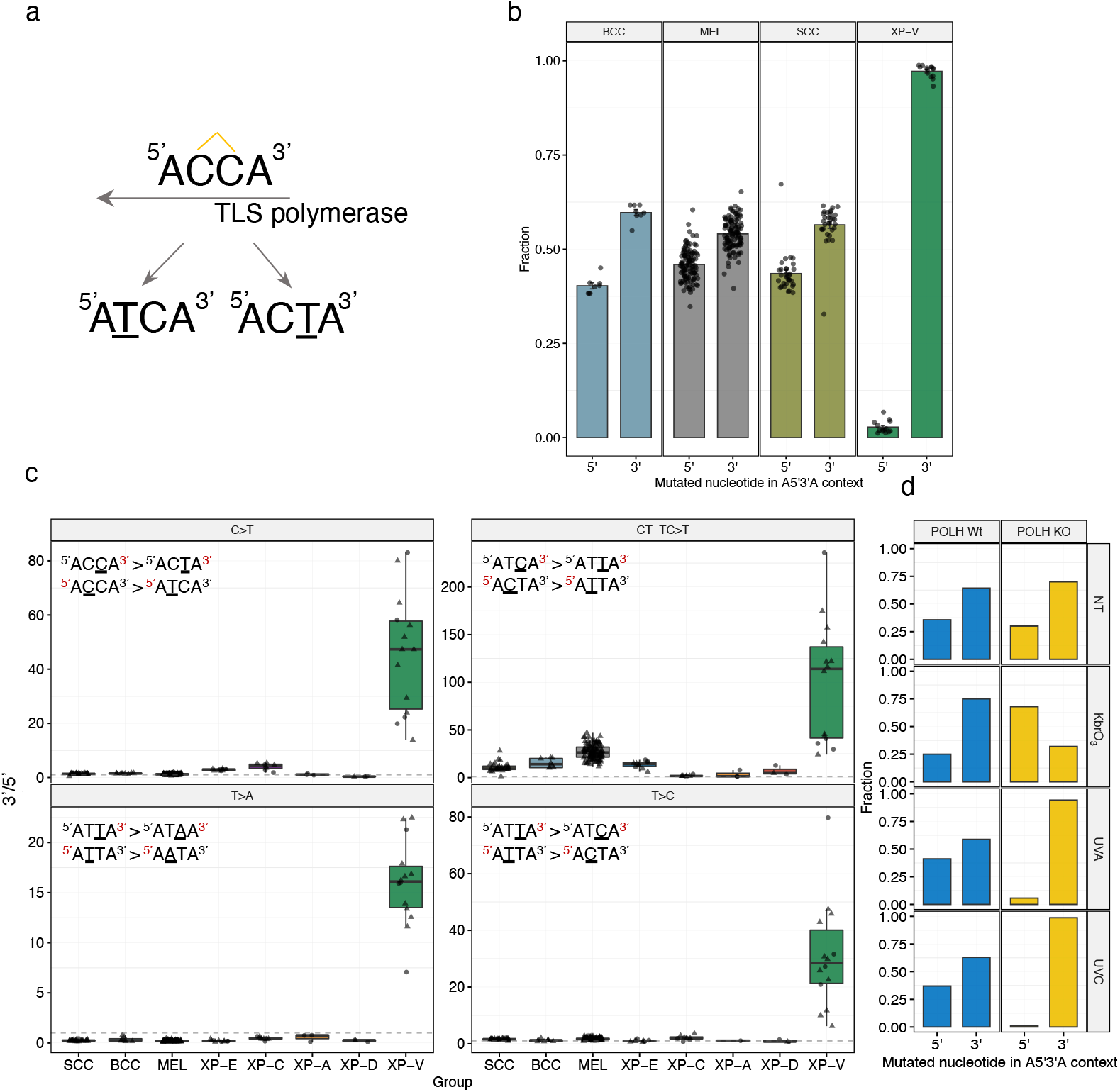
Dimer translesion bias in XP-V skin cancers. **a** Schematic representation of the putative CC photodimer in ACCA context and resulting mutations analyzed in the panel b. **b** Fraction of C>T mutations from 5’ and 3’ cytosines of the dimer in the ^5’^ACCA^3’^ context per group of tumors. **c** “Dimer translesion bias” for different sequence contexts per group of tumors. Comparison of C>T mutation frequency in CT and TC pyrimidine dimers was performed after normalization to the number of such contexts in the genome (upper right panel). **d** Fraction of C>T mutations from 5’ and 3’ cytosines of the dimer in the ^5’^ACCA^3’^ context in the RPE-1 *POLH*-wt and *POLH*-KO clones.

These results demonstrate that mutations at pyrimidine dimers in XP-V occur predominantly at the 3’ nucleotide, which might be associated with the error-prone activity of the inserter polymerase which replaces polymerase η, and modulate the mutational profile of C>T substitutions. *POLH*-KO cells treated with UV-C conversely demonstrated a very strong bias in CC pyrimidine dimers towards mutations at 3’C (99%) (**Fig. 6d**).

### Mutation properties of XP groups modulate protein-damaging effects of mutagenesis

High mutation rates in cells increase cancer risk and intensify tumor evolution, while the topography of mutagenesis and mutation signatures can impact the probability of damaging or driver mutations.^27^ In our dataset of skin cancers, the number of oncogenic mutations in the cancer genome was strongly correlated with the total mutation burden (**Fig. 7a**).

**Figure 7.**
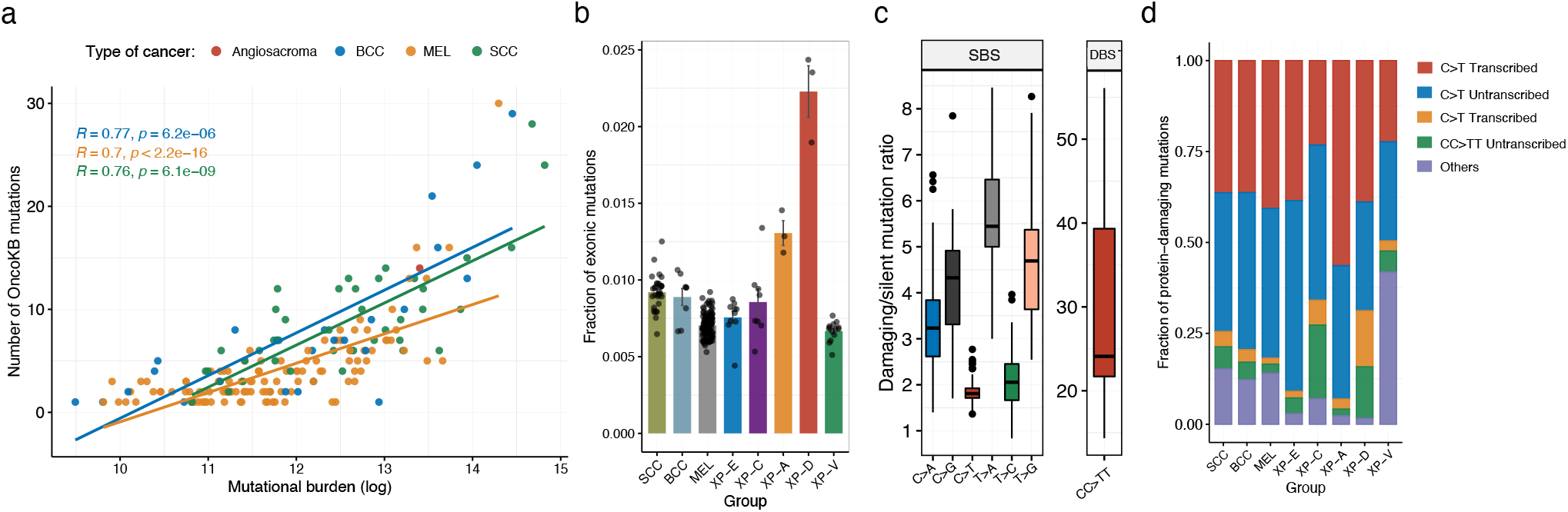
Protein-damaging effect of mutation contexts. **a** Correlations between tumor mutation burden and number of oncogenic and likely oncogenic mutations in the studied skin cancer samples according to the OncoKB database. **b** Mean fraction of exonic mutations from all the mutations per sample. **c** Protein-damaging/silent mutation ratio per substitution type in our skin cancer cohort. Damaging mutations - all non-silent exonic (missense, truncating) and splice site mutations. **d** Mean fraction of protein-damaging mutations originating from the main mutation classes split by gene strand per group.

Active DNA repair in open chromatin regions decreases the accumulation of mutations in the early replicating gene-rich regions of cancer genomes (**Fig. 2a**). We estimated a fraction of mutations per genome falling in the exonic regions across the studied skin cancer groups and found in XP-A and XP-D tumors a significant enrichment of exonic mutations in comparison with the other groups (**Fig. 7b**). The effect was caused by the redistribution of mutations from late to early RT regions of a genome (**Fig. 2a**).

C>T transitions, which are the most prevalent UV mutations, have relatively low protein-damaging effect in the human genome and their damaging/silent mutation ratio is 1.8, while other types of mutations, such as C:G>A:T transversions or CC>TT DBS are more damaging with a damaging/silent mutation ratio of 3.4 and 29.5, respectively (**Fig. 7c**). To better understand how the NER deficiency modulates the protein-damaging effect of UV irradiation we grouped protein-damaging mutations into 5 categories: C>T mutations on the transcribed and untranscribed strand, CC>TT double base substitutions on the transcribed and untranscribed strands, and other SBSs (**Fig. 7d**). The largest fraction of protein-damaging mutations was accounted for by C>T substitutions in all cancer groups except XP-V where other mutation classes play a more important role.

Contribution of damaging C>T mutations from transcribed and untranscribed strands of genes (measured as untranscribed/transcribed ratio) differed between groups. It was balanced between strands in sporadic skin cancers (1.02-fold); at the same time the majority of damaging mutations in GG-NER deficient XP-E and XP-C groups were attributed to the untranscribed strand (1.36 and 1.82-fold, respectively), while in GG- and TC-NER deficient XP-D and XP-A groups - to the transcribed strand (0.77-fold and 0.65-fold, respectively, **Fig. 7d**). These results can be explained by the fact that UV-induced C>T SBS, which originate from the lesions on the transcribed strand, are 1.88-fold more protein-damaging as compared to the untranscribed strand of genes; thereafter, active lesion removal by TC-NER from the transcribed strand of genes results not only in reduction of a total number of mutations from UV lesions, but is particularly important for the reduction of the burden of protein-damaging mutations.

## DISCUSSION

Our results indicate that XP skin cancers deficient in GG-NER (XP-C, XP-E) or polymerase η (XP-V) harbor 3.6-fold more mutations than sporadic skin cancers, on average. The mutation profiles in all XP groups were dominated by C>T mutations in pyrimidine dimers; however, they differed from sporadic skin cancers and each other. These differences can be partially explained by the increased contribution to mutagenesis of early replicating GC-rich regions in XP groups with significantly impaired NER (XP-C, XP-A, XP-D). Mutational differences in XP-E might be further explained by the important role of XPE (DDB2) protein in removal of CPD lesions rather than of 6-4PP lesions, which can be recognized by XPC directly^28^. Thus, observed distinct mutational profiles in XP-E and XP-C might be associated with the different relative contributions of CPD and 6-4PP lesions to mutagenesis. Moreover, CC>TT double base substitutions, a characteristic feature of skin cancers, which is particularly enriched in XP-C (but depleted in XP-E), could be associated with 6-4PP photolesions. In addition, CC>TT DBS were not strongly enriched or depleted in XP-V, which might suggest that the occurrence of these mutations does not depend exclusively on polymerase η.

The current knowledge about TLS across UV lesions posits that CPD are bypassed by polymerase η alone, while bulkier 6-4PP require two TLS polymerases, one of which performs insertion and the other an extension^29^. In XP-V cancer genomes, we demonstrated a striking increase in the mutagenicity of the 3’ nucleotide of the pyrimidine dimer, a phenomenon observed before using lacZ mutational reporter gene in polymerase η - deficient mice.^30^ This might be explained by the model where a CPD in the absence of polymerase η is bypassed in two steps instead of one, by an error-prone inserter polymerase followed by an error-free extender polymerase. We propose to name the effect of differential mutagenicity of 3’ and 5’ nucleotides in the intrastrand crosslinked DNA dimers as “dimer translesion bias”. In this study, we presented an illustrative example of XP-V, where a dimer translesion bias significantly alters the relatively conserved UV-induced mutation profiles for C>T mutations and drives the mutator phenotype of XP-V tumors and *POLH*-KO cell line.

It is well known that the local mutation rate in cancers and in the germline strongly correlates with epigenetic features of the genomic regions^31,32^. The most striking associations were observed for replication timing, chromatin accessibility, active and non-active topologically-associated domains, and some chromatin marks such as H3K9me3, H3K27me3, and H3K9me2^32^. In NER-deficient XP-A and XP-D tumors, we observed weak heterogeneity of the mutation frequency across the genome depending on the chromatin status. This finding demonstrates that the decreased mutagenesis in sporadic skin cancers in large open chromatin regions is driven by their accessibility to NER.

We observed that intergenic genomic regions and untranscribed strands of genes in GG-NER-deficient XP-C samples had a decreased mutation load in early-replicating genomic regions, which are enriched in genes with high levels of transcription. We have shown that it can partially be explained by the activity of TC-NER up to 50 Kb beyond the annotated TES and propose to name this phenomenon “extended TC-NER”. In the early-replicating genomic regions, genes are densely located and transcribed colinearly or in opposite orientations. Extended TC-NER can contribute substantially to the DNA repair independently of GG-NER, thus lowering mutation load in intergenic regions and on the untranscribed strands of closely located or overlapping genes.

Properties of C:G>A:T and T:A>A:T mutations, characteristic of XP-V skin tumors, such as the specific dinucleotide context (TA/G), strong transcription bias, and correlation with the number of C>T UV-induced mutations, enabled us to speculate that these mutations are associated with lesions that directly or indirectly induced by UV. C:G>A:T mutations are unlikely to be associated with 7,8-dihydro-8-oxoguanine (8oxoG) DNA adducts as mutation signatures of 8oxoG (COSMIC signatures SBS18 and SBS36) have different 3-nt context (lack of 5’ thymine specificity) and do not demonstrate transcriptional bias (https://cancer.sanger.ac.uk/signatures/sbs/). Furthermore, rare photolesions in a TA context have been previously reported, and their chemical structure was described as Thymidylyl-(3′–5′)-Deoxyadenosine “TA photoproducts”^33–35^. More recently, a study dedicated to the discovery of atypical photolesions reported rare mutagenic TA photoproducts and, to a lesser extent, TG photoproducts as the most associated with UV-irradiation after CPD and 6-4PP photolesions^36^. The high mutagenesis in TA/G contexts in XP-V tumors uncovers a critical and non-redundant function of polymerase η in the error-free bypass of highly mutagenic but poorly studied DNA lesions, probably induced by UV. The WGS analysis of the RPE-1 *POLH*-KO clones confirmed that T<u>G</u>>T<u>T</u> and T<u>A</u>>T<u>T</u> mutations are greatly increased after UV-A and UV-C exposure, and have shown that they are most prevalent after UV-A. However, after KbrO_3_ treatment which induces reactive oxygen species, in *POLH*-KO cells SBS mutational pattern did not change as compared to the wild type. Presence of TG>VA DBS in XP-V tumors and in *POLH*-KO cells after UV-exposures further suggests that their origin might depend on thymine-guanine dimers.

Another important peculiarity of the XP-V mutation profile is a high fraction of mutations (on average 15%) originating from a TT dinucleotide (T<u>T</u>>T<u>A</u>/<u>C</u>). It is known that the majority of CPD lesions occur in a TT context^37^ and polymerase η is the main polymerase to bypass them in an error-free manner. In the absence of polymerase η, other TLS polymerases perform bypass of TT lesions introducing more errors. Even though TT CPD represents near 50% of UV-induced lesions, the proportion of T<u>T</u>>T<u>A</u>/<u>C</u> mutations is only 15% in XP-V. This means that even in the absence of polymerase η, TT CPD is not a highly mutagenic lesion, probably because other replacing TLS polymerases insert predominantly adenines opposite to the lesion following the “A rule”^38,39^.

In XP skin cancers, UV irradiation results in different mutation profiles and topography of mutagenesis, which are associated with a variation in the probability of protein-damaging and oncogenic mutations. We revealed three main factors contributing to heterogeneity in the proportion of protein-damaging mutations in XP skin cancers, which were associated with the differences in mutation profiles (fractions of transversions and DBS), the activity of TC-NER, and mutation distribution between open and closed chromatin.

The observed differences in mutation burden and mutation profiles might also reflect the differences in clinical manifestations between XP groups. XP-A and XP-D patients show severe sunburn reactions and are diagnosed early. Thereafter, those patients are rarely exposed to UV and have rather few tumors. This might partly explain rather low TMB in XP-A and XP-D skin tumors. In tumor-prone XP-C, XP-E, and XP-V groups, the amount and mode of exposure to UV again may be different. XP-C patients do not experience sunburns but develop other skin symptoms resulting in an early diagnosis and sun protection. XP-E and XP-V patients, on the contrary, do not have any symptoms till their 20s or 30s, by which time they may have had a lot of sun exposure, and in subsequent decades, they develop many skin tumors. This might be in line with the observation that mutational profiles in XP-E and XP-V in some features resemble sporadic cancers, while XP-C is the most different.

Overall, our analysis of rare skin cancers with deficient NER or translesion DNA synthesis has revealed how the absence of different NER components modulates mutation burden, profiles, and topography of mutagenesis after UV irradiation. We have attempted to provide mechanistic explanations for the mutation consequences of DDB2 (XPE) loss in the XP-E group and the polymerase η deficiency in the XP-V group for UV mutagenesis in skin cancer. Further mutation studies on experimental cell lines from XP patients can extend our knowledge of the role of major and rare photoproducts in skin cancer pathogenesis and biological mechanisms supporting genome stability.

## ACKNOWLEDGMENTS

S.I.N. was supported by grant Foundation ARC 2017, Foundation Gustave Roussy, and the The French National Cancer Institute - RPT21145LLA. P.L.K and S.I.N. were supported by grant from the Foundation ARC-ARCPGA12019120001055_1578 (P.L.K. and S.N.). This work was also supported by Prism – National Precision Medicine Center in Oncology funded by the France 2030 program and the French National Research Agency (ANR) under grant number ANR-18-IBHU-0002. The authors are very thankful to Xiaole Xu (BGI) for the management of sequencing.

## AUTHOR CONTRIBUTION

S.I.N. and A.A.Y. designed the study. A.A.Y. performed the data analysis and prepared figures. A.A.Y. and S.I.N. drafted the manuscript. A.S., A.L., and C.F.M.M commented on the manuscript. F.R. handled biopsies, performed QC of the samples and DNA extraction. F.R. performed cell line experiments. P.L. participated in the DNA extraction and sample handling. K.G. participated in the data analysis. I.P. and L.P. performed data preprocessing. T.B.P., C.F.M.M., H.F., A.L., C.N., P.L.K, and A.S. collected the samples.

## DATA AVAILABILITY

Experimental data generated in this study have been deposited in the European Genome-phenome Archive (EGA) under accession XXX.

## MATERIALS AND METHODS

### Studied samples

The samples were collected from patients with a confirmed XP diagnosis. Informed signed consents were obtained from patients and/or their parents per the Declaration of Helsinki and the French law. This study was approved by the French Agency of Biomedicine (Paris, France), the Ethics Committee from the CPP of the University Hospital of Bordeaux (Bordeaux, France), the Institutional Review Board of Gustave Roussy (CSET: 2018-2820; Gustave Roussy, Villejuif, France), the Research Ethics Committee of Guy’s and St Thomas’ Foundation Trust, London (reference 12/LO/0325), and the CONEP (Brazil), Number CAAE 48347515.3.0000.5467.

The tumor samples were collected from patients during surgery. The tumors were stored in liquid nitrogen or allprotect tissue reagent, and 16 in FFPE. Normal control samples were represented by blood (4 patients), saliva (7 patients), fresh skin (2 patients), or FFPE (6 patients). DNA from non-FFPE tissues was extracted using AllPrep DNA/RNA/miRNA Universal Kit (Cat. No. / ID: 80224, Qiagen) according to the manufacturer’s instructions. DNA from FFPE blocks was extracted after examination and dissection by a pathologist. Tumor DNA was extracted from parts of FFPE containing a high fraction of tumor cells using Maxwell® RSC DNA FFPE Kit (Catalog number: AS1450, Promega) according to the manufacturer’s instructions. Non-tumoral DNA was extracted from FFPE blocks that did not contain tumor cells if available, or from parts of tumor cell-containing FFPE blocks free from tumor cells. DNA quantity and quality were assessed using the NanoDrop-ND-1000 (Nanodrop Technologies).

### Genome sequencing and variant calling

The genomes were sequenced using BGISEQ-500 in BGI (Shenzhen) according to the manufacturer’s protocols to the mean coverage after deduplication equal to 40X for tumor and 30X for normal DNA (100 bp paired-end reads). Reads were mapped using BWA-MEM^40,41^ (v0.7.12) software to the GRCh37 human reference genome, and then we used the standard GATK best practice pipeline^42^ to process the samples and call somatic and germline genetic variants. PCR duplicates were removed, and the base quality score recalibrated using GATK^43^ (v4.0.10.1), MarkDuplicates, and BaseRecalibrator tools. Somatic variants were called and filtered using GATK tools Mutect2, FilterMutectCalls, and FilterByOrientationBias and annotated with oncotator^44^ (v1.9.9.0). SCNAs calling was done with FACETS^45^ (v 0.5.14). Quality controls of FASTQ files and mapping were done with FASTQC^46^ (v0.11.7), samtools^47,48^ (v1.9), GATK HSmetrics, and MultiQC^49^ (v1.5). All processing steps were combined in a pipeline built with snakemake^50^ (v5.4.0).

### Filtration of somatic variants in tumor samples

Only PASS-filtered somatic variants supported by at least one read from each strand and at least three reads in total with variant allele frequency higher than 0.05 and POPAF filter > 5 (negative log 10 population allele frequencies of alt alleles; probability of the mutation to be a germline polymorphism) were used for the analysis. Additionally, all used VCF files were filtered based on the alignability map of the human genome from the UCSC browser (https://genome.ucsc.edu/cgi-bin/hgFileUi?db=hg19&g=wgEncodeMapability) with the length of K-mer equal to 75 bp (wgEncodeCrgMapabilityAlign75mer, mutations overlapped regions with score <1 were filtered out) and UCSC Browser blacklisted regions (Duke and DAC).

To filter out the FFPE artefacts, we employed Support Vector Machine-based (SVM) methodology with the e1071 R library.^51^ For each sample separately, each variant in the prefiltered VCF file (the same filters as for the fresh non-FFPE samples) was annotated with additional quality information specific for the alternative allele from the BAM file using bam-readcount utility^52^. This additional BAM-derived information in the form of a table was merged with the quality annotations from the VCF file (VCF was parsed into a table with vcf2tsv from vcflib library^53^) which included CONTQ (Phred-scaled qualities that alt allele are not due to contamination), SEQQ (Phred-scaled quality that alt alleles are not sequencing errors), STRANDQ (Phred-scaled quality of strand bias artifact), TLOD (Log 10 likelihood ratio score of variant existing versus not existing). The typically UV-induced double base substitutions (CC:GG>TT:AA) were considered true positive variants, while abundant FFPE artefacts TG:CA>CA:TG were considered false positive variants during the training of SVM. To tune the SVM parameters we subset 25% of the TG:CA>CA:TG and CC:GG>TT:AA variants and run tune() command (cost=c(0.001,0.01,0.1, 1,5,10,100)). Then the best tuning parameters for the model were chosen (tune.out$best.model) and applied to the training dataset of 50% of the TG:CA>CA:TG and CC:GG>TT:AA variants using svm() command with 10 k-fold cross validations (cross=10) and probabilistic assignment of the classification (type=“C-classification”, probability = TRUE, scale=T) to build the SVM classification model. Finally, the SVM classification model was applied to the whole dataset of variants to classify them as true positive in a probabilistic manner (command predict(), probability = TRUE). We extracted for the downstream analysis only the variants with a probability of being true positive > 0.95.

### Mutation spectrum, MDS, and comparison with known signatures

To convert the VCF files into a catalog of mutation matrices, we used SigProfilerMatrixGenerator v.1.0 software^54^. Before the profiling, VCF files were split into separate files with single base substitutions and other variants to avoid splitting double base substitutions into single base substitutions by the software. To construct the multidimensional scaling plots (MDS), we computed pairwise Cosine similarity distance between all the samples using MutationalPatterns R package^55,56^ (cos_sim_matrix()) and then processed the matrix of distances between the samples in the prcomp() function in R.

To understand whether known UV signatures can explain the mutational profiles of XP and sporadic datasets, we extracted four SBS mutational signatures previously associated with UV irradiation (SBS 7a,b,c,d) from the COSMIC database^57^ (V3.2, https://cancer.sanger.ac.uk/signatures/sbs/) and then reconstructed observed mutational profiles of the studied samples using these four UV-associated mutational signatures (fit_to_signatures(), MutationalPatterns R package). The Cosine dissimilarity of the observed and reconstructed mutational profiles was calculated for each sample as 1-Cosine distance. The procedure was performed separately for all 96 trinucleotide mutational contexts and only 12 mutational contexts of the UV-induced spectra (N<u>C</u>Y>N<u>T</u>Y or Y<u>C</u>N>Y<u>T</u>N).

### Replication timing, TADs, epigenetic marks, and mutational load along the genome

We used Repli-Seq data from 11 cell lines^58^ (BG02, BJ, GM0699, HeLa, HEPG2, HUVEC, IMR90, K562, MCF7, NHEK, SK-N-SH) to identify conservative replication timing regions. For each 1-kb region, we calculated weighted mean replication timing and then its standard deviation between all the cell lines and removed all the regions with a standard deviation higher than 15. For the rest of consistent regions across different cell lines, we calculated the mean values and used them during analysis. The genome was divided into five or eight bins according to the replication timing values, and mutation density was calculated for each bin, adjusting for trinucleotide contexts. Additionally, we computed the dependence of mutation density on replication timing separately for intergenic and genic regions (splitting mutations on the transcribed and untranscribed strands).

The genomic location of the 1MB borders between topologically-associated domains (TADs) was downloaded from the recent publication exploring mutation rate dependency on TAD structures^19^. The border regions were spitted into 1-kb intervals and separated into four bins (two for active and two for inactive TADs). Then the fraction of mutations per each sample fallen into each bin was calculated, adjusting for the trinucleotide composition. A similar procedure was performed for the consensus chromatin states of the genome from the same publication.

To calculate the slopes of the mutation load over replication timing (or other epigenetic marks) bins per sample, the logarithm of the normalized fraction of mutations in each bin was fitted into a linear model (lm()) with the number of each bin (1 to 8).

To investigate the relationships between mutation density and intensity of various epigenetic marks (DNase, H3K36me3, H3K27ac, H3K4me1, H3K27me3, H3K9me3, methylation level from whole genome bisulfite sequencing), we downloaded bigwig files of the Roadmap Epigenomics Project^25^ and converted them to wig and then bed files (tissue E058, keratinocyte). The mean intensity of each mark was calculated for 1-kb non-overlapping windows across autosomes with BEDOPS v2.4.37 (bedmap) software.^59^ The mark intensities were normalized to the 1−100 range, and we used only genomic windows with high alignability (equal to 1) along at least 90% of a window. For each window, we split mark intensities into 5 bins (cut2() function in R) and calculated the trinucleotide-adjusted fraction of mutations per sample per bin for each mark separately for intergenic regions, transcribed and untranscribed strands of genes.

To assess the mutation load distribution along the genome between groups of samples and irrespective of the epigenetic features, we split the genome into 1MB-long nonoverlapped intervals and excluded all the intervals with a mappability score less than 1 over 80% of the interval. For the resulting dataset of 2684 intervals, we calculated the mutation density of C>T substitutions in each interval per sample (with at least 50000 mutations) and then normalized the mutation density. Finally, the principal component analysis was performed on the resulting matrix.

### Transcriptional bias and XR-seq

Transcriptional strand bias (TRB) was quantified for each sample based on the stranded mutation matrixes generated by SigProfilerMatrixGenerator.^54^ We computed inequality between mutations from pyrimidines (C > A/T/G; T > A/C/G) to mutations from purines (G > A/C/T; A > C/G/T) for genes located on the sense and antisense strands of DNA relative to the reference human genome.

To compute TRB between genes expressed with different levels, we used RPKM values of RNA-seq from Epigenetic Roadmap Project^25^ represented by keratinocytes (E058) and only samples represented by BCC and cSCC. For each gene, mutations were separated as located on transcribed or untranscribed strands, and genes were divided into six bins by the level of expression.

Following the hypothesis that cytosine-containing DNA lesions caused the majority of mutations, we were also able to compute strand-specific mutation densities around transcription end sites (TESs), and transcription start sites (TSSs). Transcribed and untranscribed strands of genes and adjacent to TES/TSS intergenic regions were treated separately. TESs/TSSs of all annotated genes (GENECODE^60^ v38) were retrieved using BEDTools v2.30.0^61^, and then regions located ±50 kb of TESs/TSSs were split into 1-kb intervals. The 1-kb intervals that overlapped with other intergenic or genic intervals (represented mainly by overlapped or closely located genes) were removed for this analysis, and the rest were aggregated into 10 bins. We then separately calculated the trinucleotide context-adjusted fraction of mutations per bin per sample for transcribed and untranscribed strands.

XR-seq profiles for XP-C cell lines (XP4PA-SV-EB, GM15983) and nascent RNA-seq data from the HeLa cell line were downloaded from the previous works of Hu et al. 2015^22^ and Barbieri et al. 2020^23^, respectively. The mean intensity of tracks was calculated for binned 1-kb intervals along the genome and ±50 kb around the TESs.

### Dimer translesion bias

To calculate the relative amount of mutations arising from 5’ and 3’ sides of pyrimidine dimers, we extracted mutations from C>T located in the ACCT context, mutations T>C/A located in the ATTA context, and calculated the ratio of such mutations originating from 3’ C/T to 5’C/T separately for each mutation type with the corresponding 4-nucleotide context. Additionally, we calculated the ratio between the number of AT<u>C</u>A > AT<u>T</u>A and A<u>C</u>TA > A<u>T</u>TA mutations per sample, adjusting for the different fractions of AT<u>C</u>A and A<u>C</u>TA four-nucleotides.

### Protein-damaging effects of mutagenesis

To assess the protein-damaging effect of different substitutions, we annotated the VCF files using oncotator^44^ software and classified exonic mutations into protein-damaging (missense, nonsense, splice-site) and silent. For the C>T and CC>TT mutations, we separated them by strands and calculated the protein-damaging effect separately for the transcribed and untranscribed strands. The number of putative oncogenic drivers per sample was calculated using the OncoKb^62^ database (oncogenic and likely oncogenic events).

### Cell culture

RPE-1 *TP53*-KO cell line is obtained as a gift from Dr. Olivier Gavet lab. RPE-1 *POLH*-wt and RPE1 *POLH*-KO cell lines were cultured in DMEM/F-12 (gibco; life technologies, Ref: 11320033) at 37 °C in a humidified atmosphere containing 5% CO2., supplemented with 10% (v/v) fetal bovine serum (FBS; NB-26-00009).

### Generating *POLH*-KO cell line

*POLH*-KO cell lines are obtained from Synthego company. sgRNA was used to generate the *POLH*-KO cell line. Homozygote knock out was verified by sanger sequencing showing 4 nucleotide insertion.

### UV exposure

Cells were irradiated with 10 J/m^2^ UV-C (200-280 nm) or UV-A (320-400 nm) for 4 sequential exposures both for *POLH*-KO and *POLH*-wt cell lines. Irradiation were performed every 4 days.

### KbrO_3_ treatment

IC50 values for KbrO_3_ was identified as following protocol. 5,000 Cells per each well were plated and grown for 24 hours in 96-well plates. Cells were treated in serial diluted concentrations of KbrO_3_ (500mM-10uM). Treatment was last for 96 hours. After 4 days, density of cells in each well was quantified using Methylene Blue staining. In the first step, cells (wells) were washed with PBS 1X. Then 100μl absolute methanol is added to each well and plate was incubated for 1 hour at room temperature. Then the wells were let to be dried and 100 μl methylene blue solution (concentration 1gr/L) was added to each well and followed by 1-hour incubation at room temperature. Following the staining step, wells were rinsed with water for 2 times and then let the wells to dry. Washing step followed by solubilization of stain by adding 200 μl HCL (0.1N) in each well and incubation at 60°C for 30 minutes. In the last step O.D of each well was measured at 630 nm using BMG FLUOstar OPTIMA plate reader.

*POLH*-wt and *POLH*-KO cells were treated for 8 weeks using 300uM KbrO_3_. Treatment was refreshed every 48 hours.

### Single-cell cloning and DNA extraction

Single cell sorting was performed by Flow Cytometry Cell Sorting (FACS) in P96-well plate upon the completion of treatment period and reaching the sufficient number of cells. 5-6 hours after sorting the wells were monitored to confirm the presence of a single cell in the well. Three clones per condition were randomly selected to pass to P24-well plate to propagate the cells and extract DNA 18-21 days after cell sorting. Genomic DNA was extracted using the Qiamp DNA mini kit (QIAGEN) according to the manufacturer’s instructions.

### Sequencing and bioinformatic analysis of the cell line experiments

The RPE-1 clonal cell populations were sequenced in BGI, Shenzhen (15x coverage, BGISEQ-500 instrument) and bioinformatically processed in the similar way with tumor samples. The nontreated samples were used as “normal” and treated as “tumor” during GATK mutect2 calling of somatic mutations (and vice versa). Then we removed all the nonunique mutations between the clones (module *bcftools iseq*) as well as supported by less than 3 reads in total and at least one read from each strand. Finally, only strictly clonal mutations with VAF > 0.3 were used for the analysis.

## TABLES

**Supplementary Table 1. Xeroderma Pigmentosum tumors used in the analysis**.

## SUPPLEMENTARY FIGURE LEGENDS

**Supplementary Figure 1**. Trinucleotide-context mutation profiles of SBS for each tumor from XP patients.

**Supplementary Figure 2**. Fractions of C>T mutations from pyrimidine dimers in intergenic regions (INT, grey color), on the untranscribed (NTR, red color) and transcribed (TR, blue color) DNA strands of gene regions grouped in 5 equal size bins by replication timing (RT) for XP groups and sporadic skin cancers.

**Supplementary Figure 3**. Transcriptional bias in the TES-centered 100kb region (binned by 10kb intervals).

**Supplementary Figure 4**. Correlations between different types of substitutions in specific contexts in XP-V tumors.

**Supplementary Figure 5**. Scheme of the mutation accumulation experiment with *POLH* KO and *POLH* wt cell lines.

**Supplementary Figure 6**. Double base substitution (DBS) profiles of XP and sporadic skin tumors from fresh-frozen samples (upper panel) and RPE-1 mutation accumulation experiment (lower panel). Only fraction from 0 to 0.3 is shown.

