## Supplementary Figures 1-6 for "Analysis of Skin Cancers from Xeroderma Pigmentosum Patients Reveals Heterogeneous UV-Induced Mutational Profiles Shaped by DNA Repair"

### Supplementary Figure 1

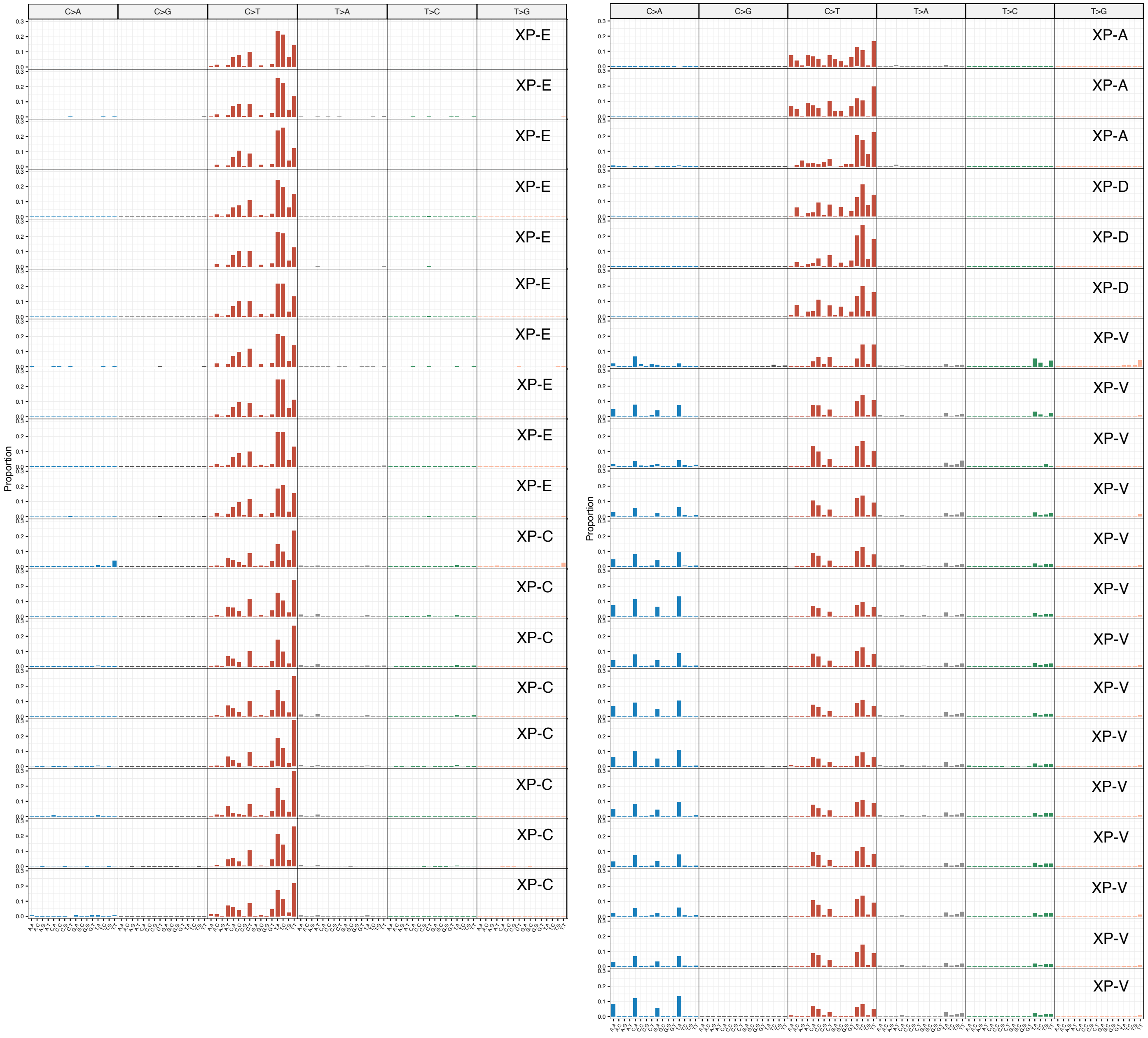


### Supplementary Figure 2

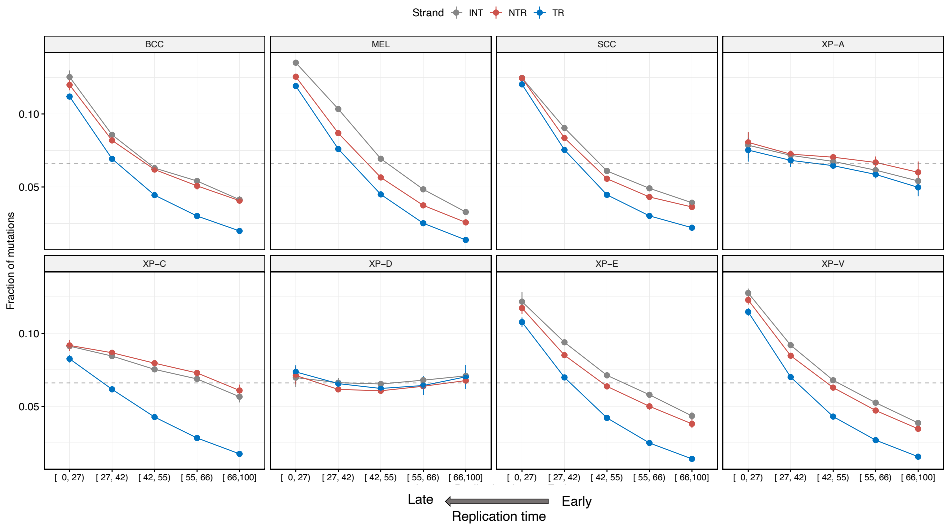

#### Supplementary Figure 3

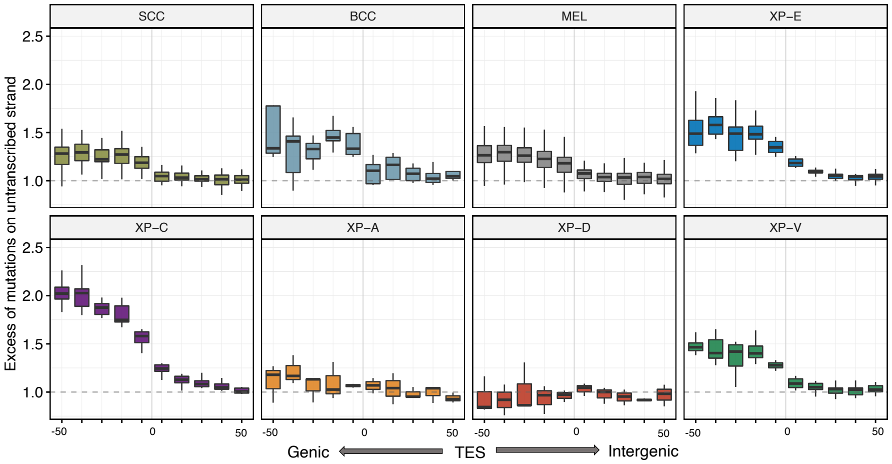


#### Supplementary Figure 4

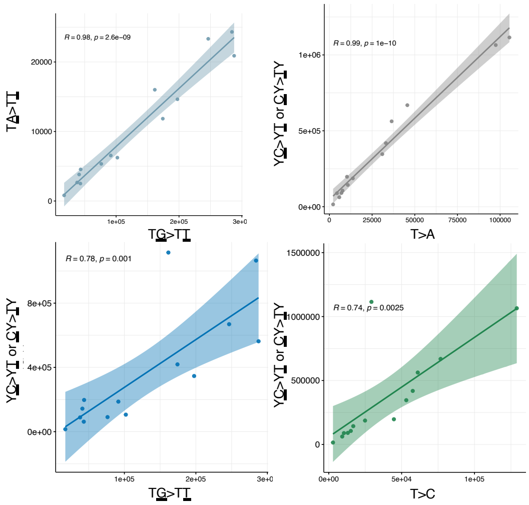


#### Supplementary Figure 5

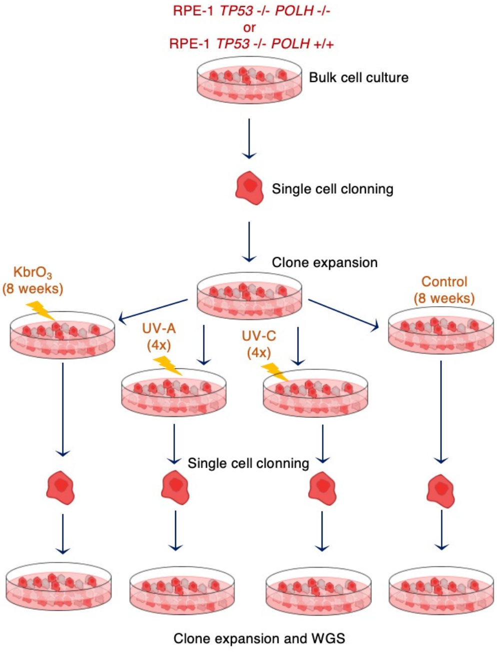

### Supplementary Figure 6

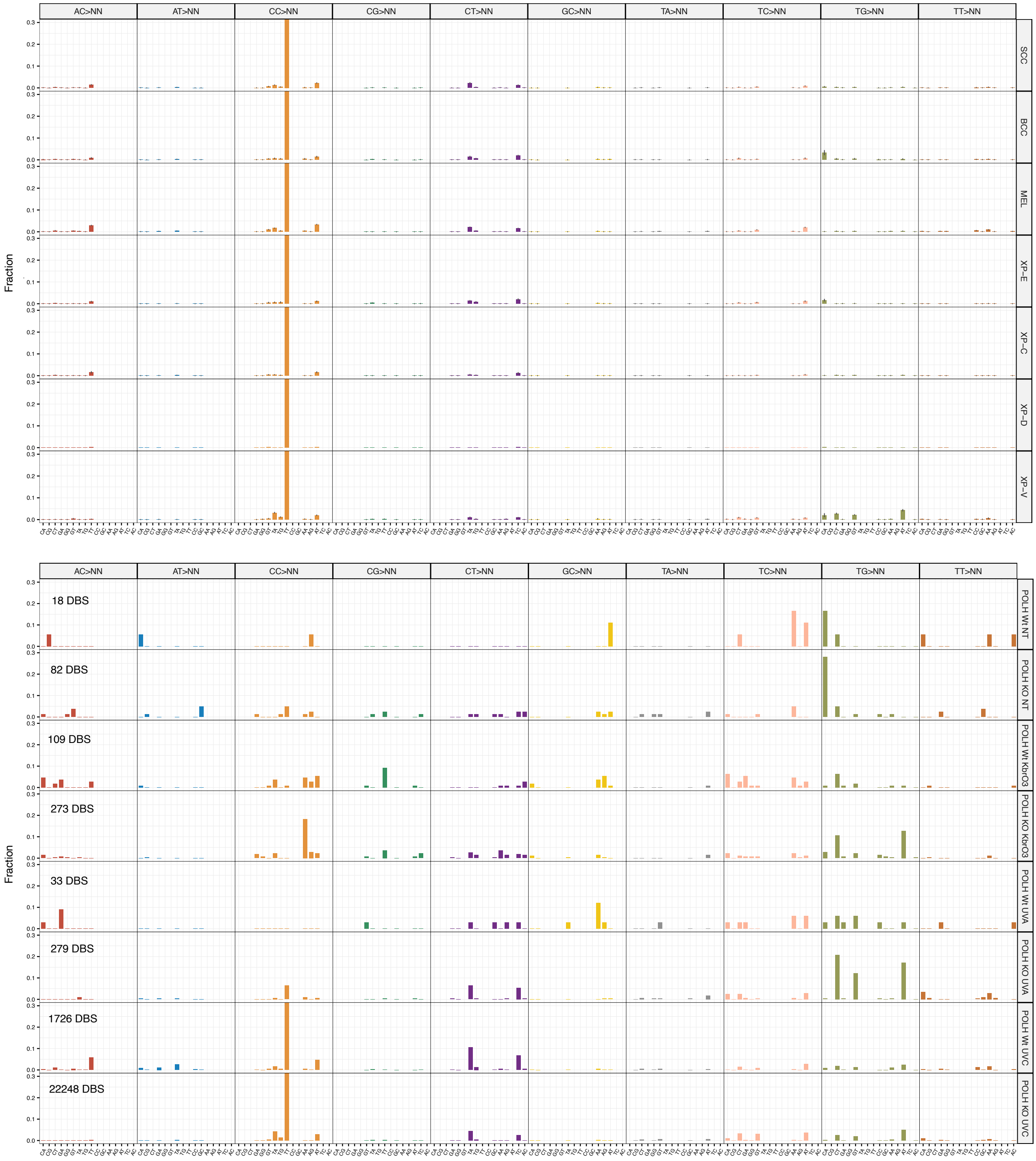
